# Effects of endophyte infected tall fescue seed and red clover isoflavones on rumen microbial populations, fiber fermentation, and volatile fatty acids *in vitro*

**DOI:** 10.1101/377085

**Authors:** Emily A. Melchior, J. Travis Mulliniks, Jason K. Smith, Gary E. Bates, Liesel G. Schneider, Zachary D. McFarlane, Michael D. Flythe, James L. Klotz, Jack P. Goodman, Huihua Ji, Phillip R. Myer

**Affiliations:** Department of Animal Sciences, University of Tennessee, Knoxville, TN, USA; Department of Plant Sciences, University of Tennessee, Knoxville, TN, USA; USDA-ARS, Forage-Animal Production Research Unit, Lexington, KY, USA; Department of Plant and Soil Sciences, University of Kentucky, Lexington, KY, USA; Kentucky Tobacco Research & Development Center, University of Kentucky, Lexington, KY, USA

**Author notes:** Corresponding Author: email - (PRM). West Central Research and Extension Center, University of Nebraska-Lincoln, North Platte, NE 69101, USA. California Polytechnic State University Department of Animal Sciences, San Luis Obispo, CA 93407, USA.

**Keywords:** rumen, cattle, tall fescue, clover, isoflavones

## Abstract

Negative impacts of endophyte-infected *Lolium arundinaceum (Darbyshire)* (tall fescue) are responsible for over $2 billion in losses to livestock producers annually. While the influence of endophyte-infected tall fescue has been studied for decades, mitigation methods have not been clearly elucidated. Isoflavones found in *Trifolium pretense* (red clover) have been the subject of recent research regarding tall fescue toxicosis mitigation. Therefore, the aim of this study was to determine the effect of ergovaline and red clover isoflavones on rumen microbial populations, fiber degradation, and volatile fatty acids (VFA) in an *in vitro* system. Using a dose of 1.10 mg × L^−1^, endophyte-infected or endophyte-free tall fescue seed was added to ANKOM fiber bags with or without 2.19 mg of isoflavones in the form of a control, powder, or pulverized tablet, resulting in a 2 × 3 factorial arrangements of treatments. Measurements of pH, VFA, bacterial taxa, as well as the disappearance of neutral detergent fiber (aNDF), acid detergent fiber (ADF), and crude protein (CP) were taken after 48 h of incubation. aNDF disappearance values were significantly altered by seed type *(P* = 0.003) and isoflavone treatment (*P* = 0.005), and ADF disappearance values were significantly different in a seed x isoflavone treatment interaction (*P* ≤ 0.05). A seed x isoflavone treatment interaction was also observed with respect to CP disappearance (*P* ≤ 0.05). Seventeen bacterial taxa were significantly altered by seed x isoflavone treatment interaction groups (*P* ≤ 0.05), six bacterial taxa were increased by isoflavones (*P* ≤ 0.05), and eleven bacterial taxa were altered by seed type (*P* ≤ 0.05). Due to the beneficial effect of isoflavones on tall fescue seed fiber degradation, these compounds may be viable options for mitigating fescue toxicosis. Further research should be conducted to determine physiological implications as well as microbiological changes *in vivo.*

## Introduction

Ergot alkaloid toxins found in the endophyte-infected tall fescue plant contribute to drought, heat, and disease tolerance of the plant, but are implicated in expensive production losses when grazed by livestock [1]. Concerns of the alkaloids manifest with reduced blood flow to the periphery, reduced reproductive efficiency and reductions in growth performance [2–4]. Of the alkaloids produced by endophyte-infected tall fescue, ergovaline is the most predominant of concern in the plant, as it has the most detrimental effects on livestock [5, 6]. Reducing the consumption of endophyte-infected tall fescue has proven to be a challenge for livestock producers, as it occupies roughly 15 million hectares in the United States [7], with the majority in the east and southeast regions of the country. Replacing endophyte-infected varieties with novel and endophyte-free varieties of tall fescue can abate toxicosis, but may not be a cost-effective long-term solution [8]. Alternatively, the incorporation of cool-season legumes into endophyte-infected tall fescue pastures has been shown to mitigate a portion of tall fescue toxicosis symptoms [9, 10]. Phytoestrogenic compounds found in red and white clover have been reported to improve overall growth performance of livestock grazing endophyte-infected tall fescue throughout the spring and summer months [11–13]. Various isoflavones have been targeted for use in human medicine due to their beneficial reduction in breast cancer symptoms, improvements in cardiovascular health, as well as reductions in menopausal symptoms [14–16]. As such, determining additional benefits of isoflavones, such as the improvement of fiber utilization and rumen fermentation in livestock, is timely.

Ergot alkaloid pressure is reduced in the presence of ruminal fluid [17, 18], indicating that ruminal microorganisms may be responsible for degradation of ergot alkaloids to less harmful compounds such as lysergic acid. Harlow, Goodman (19) identified several hyper-ammonia producing and tryptophan-utilizing bacteria capable of degrading ergovaline. Tryptophan is essential for ergovaline synthesis and formation of the ergoline ring structure. It has been hypothesized that bacterial species able to degrade tryptophan may also degrade ergovaline in the rumen [19, 20], proving essential for reducing the effects of tall fescue toxicosis. While a phylogenetically [21] and functionally diverse rumen microbiome is essential to the health and nutritional status of ruminant animals, feed additives or plant-secondary compounds, such as isoflavones, may further improve rumen functionality in tandem with microbial populations. Isoflavones may enhance rumen function by improving fiber degradation and by reducing several microorganisms responsible for lowered ruminal pH and increased lactate production that can disrupt normal functional microflora [22, 23].

This study examined the effect of endophyte-infected or endophyte-free tall fescue seed with or without an isoflavone source on *in vitro* rumen incubation, fiber degradation, and rumen bacterial communities. The objectives of this study were to 1) determine if isoflavones improve fiber degradation and rumen incubation of tall fescue seed *in vitro*, and 2) determine if *in vitro* rumen bacterial populations are affected by seed type or the addition of isoflavones. We hypothesized that the addition of isoflavones to endophyte-infected tall fescue seed would alter *in vitro* fermentation and rumen bacterial populations.

## Materials and Methods

### Study Design

This study was approved and carried out in accordance with the recommendations of the Institutional Animal Care and Use Committee at the University of Tennessee, Knoxville. This experiment was conducted as a randomized incomplete block design, and was blocked by each Daisy^II^ run. Eight separate runs were performed. The first seven runs included four jars to which treatments were randomly allocated, and the eighth run consisted of two jars, for a total of five replications of six treatment combinations. Treatments were arranged as a 3 × 2 factorial design that included three levels of isoflavones: (1) Promensil® 80 mg isoflavone tablet or (2) Promensil® 80 mg equivalent isoflavone powder, and (3) control, receiving no isoflavones, and two seed types: (1) endophyte-infected tall fescue seed (KY 31) or (2) endophyte-free tall fescue seed (KY 32).

For the responses of aNDF and ADF disappearance, ten fiber bags per jar were utilized as the sampling units. This yielded a total number of aNDF and ADF disappearance observations of 300. For the response of CP disappearance, five bags per jar were composited to a total of two weigh boats per jar as the sampling unit. This yielded a total number of CP disappearance observations of 60. Similarly, for the responses of ruminal pH and volatile fatty acids (VFA), two samples per jar were obtained using a 15-mL conical tube, yielding 60 ruminal pH and VFA observations.

### In vitro fiber and crude protein disappearance

The Daisy^II^ *in vitro* incubation system (ANKOM Corp., Fairport, NY) was utilized to determine the rate and extent of fiber disappearance of the endophyte-infected or endophyte-free tall fescue seed with or without the addition of isoflavones from Promensil®. Each substrate (500 ± 40 mg) was weighed into artificial fiber bags (F57 fiber bags, ANKOM Corp.), which were then heat sealed. Content included 442.5 mg of either endophyte-infected (KY 31) or endophyte-free tall fescue (KY 32) seed (dry matter basis), and 57.5 mg of Promensil® (tablet or powder, dry matter basis). The fiber bags were separated into four groups of 12 bags within a seed type x isoflavone treatment combination, including two empty bags for correction, and placed into separate upright plastic containers. A total of 1600 mL of rumen fluid was procured via aspiration from two fistulated Holstein heifers (Cherokee Farm, Knoxville, TN), and buffered using a 1:4 dilution of rumen fluid to McDougall’s artificial saliva buffer [24]. A total of 400 mL were added to each ANKOM jar with 1600 mL of buffer, and adjusted to a pH of 6.8 with CO_2_. The fiber bags and diluted/buffered rumen fluid were added to the incubation containers. Incubation then proceeded for 48 h at 39 ± 0.5°C.

Upon completion of the 48-h incubation, rumen pH was measured using a pH meter (Denver Instruments, Bohemia, NY) and 15 mL of rumen content was sampled and stored at − 80°C until further processing. All fiber bags were removed from jars, and washed under tap water before further analyses. Analysis of aNDF was obtained using α-amylase and sodium sulfite [25], and ADF was determined using sulfuric acid-based detergent [26]. Both procedures were conducted using the ANKOM200 fiber analysis system (ANKOM Corp., Fairport, NY). Crude protein was determined by total nitrogen combustion analysis (LECO Instruments, Inc., St. Joseph, MI). Difference between pre- and post-fermentation aNDF, ADF, and CP levels were used to calculate disappearance.

### Quantification of ergot alkaloids

Prior to the study, concentrations of ergovaline and its epimer ergovalinine in fescue seed were determined using HPLC with fluorescence detection as described in Aiken et al. [27] with modifications described by Koontz et al. [28]. The endophyte-infected fescue seed contained 2.94 ppm of ergovaline and ergovalinine (1.85 and 1.09 ppm, respectively), and the endophyte-free tall fescue seed contained a total of 0 ppm of ergot alkaloids. Additionally, both seed varieties tested negative for the presence of the alkaloid ergotamine and its epimer ergotaminine. Seed samples were ground through a 5-mm screen using a Wiley Mill before 0.4425g of seed (1.3865 × 10^−6^ g ergot alkaloids) was added to each ANKOM fiber bag.

### Quantification of isoflavones

Isoflavones were procured by finely grinding the product Promensil® (PharmaCare Laboratories, Warriewood, NSW 2102, Australia), an over-the-counter isoflavone supplement isolated from red clover, with mortar and pestle to pass through a 1 mm screen. Quantification of isoflavones in Promensil®, including biochanin A, formononetin, genistein and daidzein, was performed using methods similar to those previously described by Aiken et al. (2016), using LC-MS rather than UV for detection. Briefly, isoflavone extracts were prepared by adding 7 mL of 85% methanol in 0.5% acetic acid to ground samples in 50-mL conical polypropylene tubes. Samples were briefly vortexed and sonicated for 30 min at ambient temperature. Three mL of deionized water was added to each sample prior to being vortexed and centrifuged for 8 min at 2200 × g. The resulting supernatant was filtered through a 0.45 μm GHP membrane syringe filter. Extracts were then diluted, and flavone added as internal standard. One portion of each sample was analyzed as-extracted and a second portion was heated at 85°C for 5 h to hydrolyze isoflavone malonyl-glucosides to their corresponding isoflavone glucosides. Concentrations of biochanin a-malonyl-glucoside and formononetin-malonyl-glucoside were determined by difference between hydrolyzed and un-hydrolyzed portions. Isoflavone extracts were analyzed by LC-MS on a Waters Acquity UPLC coupled to a Waters Synapt G2 (q-ToF) high resolution mass spectrometer. Chromatographic separation was obtained using a Waters BEH C18 UPLC column (1.7 μm, 2.1 mm × 150 mm). The mobile phase employed a mixture of water containing 0.1% formic acid (solvent A) and acetonitrile containing 0.1% formic acid (solvent B) in a linear gradient from 20% B to 80% B at a flow rate of 0.35 mL × min^−1^. The high resolution mass spectrometer was operated in positive ion electrospray mode with a resolving power of ~14,000 and scanned from 100 to 1000 Da in 0.3 s. Leucine enkephalin was used to provide a lock mass (m/z 554.2615). Quantification of isoflavones was performed using QuanLynx software with a linear calibration curve and internal standard method. Extracted ion chromatograms with a mass window of 0.02 Da around the accurate mass of each analyte were used to calculate peak areas. A total of 2.19 mg of total isoflavones were provided per bag (0.92 mg biochanin A, 0.89 mg formononetin, 0.0038 mg genistein and 0.0016 mg daidzein), for a concentration of 1.10 mg × L^−1^ of rumen fluid and buffer.

### VFA Analysis

A subsample of rumen fluid was aliquoted from each jar for VFA analysis using HPLC, similar to the procedures previously described by Harlow et al. (2017). Briefly, samples were analyzed for concentrations of acetate, propionate, butyrate, valerate, and isovalerate/methylbutyrate (IVMB) using a Summit HPLC (Dionex; Sunnyvale, CA, USA) equipped with an anion exchange column (Aminex HP-87H; Bio-Rad, Hercules, CA, USA) and UV detector. The eluting compounds were separated isocratically with an aqueous sulfuric acid solution (5 mM). The parameters included an injection volume of 0.1 mL, flow rate of 0.4 mL × min^−1^, and column temperature of 50°C.

### DNA extraction, PCR and sequencing

DNA was extracted from ruminal fluid post-fermentation. The procedure of the DNA extraction method was similar to that previously described by Yu and Morrison [29]. After the chemical/mechanical cell lysis and isopropanol precipitation of nucleic acids, metagenomic DNA was purified with Rnase and proteinase K treatment, followed by the use of QIAamp columns from the Qiagen DNA Stool Mini Kit (Qiagen, Hilden, Germany). Genomic DNA concentration was determined using a Nanodrop 1000 spectrophotometer (Thermo Scientific, Wilmington, DE), and verified using Invitrogen Qubit fluorometer with PicoGreen (ThermoFisher Scientific, Wilmington, DE). Extractions were stored at −20°C until sequencing library preparation. Bacterial 16S rRNA genes were PCR-amplified with dual-barcoded primers targeting the V4 region, as per the protocol of Kozich [30]. Amplicons were sequenced with an Illumina MiSeq using the 250-bp paired-end kit (v.2). Sequences were denoised, taxonomically classified using Greengenes (v. 13_8) as the reference database, and clustered into 97% similarity operational taxonomic units (OTUs) with the mothur software package (v. 1.39.5), as previously described by Schloss, Westcott (31), and following the manufacturer-recommended procedure (https://www.mothur.org/wiki/MiSeq_SOP; accessed November 2017).

## Statistical Analyses

The efficiency of the incomplete block design [32] was determined using PROC OPTEX in SAS 9.4 (SAS Inst. Inc., Cary, NC). Study analyses included responses of aNDF, ADF, CP disappearance, rumen pH, VFA concentrations, and ruminal bacterial communities. Separate mixed model analyses of variance (ANOVA) were performed using the MIXED procedure of SAS to determine the fixed effects of isoflavone treatment, seed type, and seed type x isoflavone treatment interaction. The ANOVA models also included random effects of run and run x seed type x isoflavone treatment. Least square means were separated using least significant differences, and Tukey’s adjustments were made for multiple comparisons. Effects were considered significant at *P* ≤ 0.05, and tendencies declared at *P* > 0.05 and ≤ 0.10.

For rumen bacterial communities, analysis was conducted in the R environment. Alpha diversity was estimated with the Shannon index on raw OTU abundance tables. The significance of diversity differences was tested with an ANOVA. To estimate beta diversity across samples, OTUs were excluded if occurring in fewer than 10% of the samples with a count of less than three and computed Bray-Curtis indices. Beta diversity, emphasizing differences across samples, was visualized using non-metric multidimensional (NMDS) ordination. Variation in community structure was assessed with permutational multivariate analyses of variance (PERMANOVA) with treatment group as the main fixed factor and using 4,999 permutations for significance testing.

## Results

### Efficiency of Incomplete Block Design

Efficiency is a measurement of goodness of an experimental design [32]. D-efficiency is calculated based on the geometric mean of eigenvalues of the variance matrix [32]. In a perfectly balanced design, D-efficiency will equal 100%. In our study, the treatment D-efficiency was 87.6% for our allocation of treatments to runs.

### aNDF, ADF, and CP Disappearance

Endophyte-infected tall fescue seed had significantly different aNDF disappearance when compared to endophyte-free tall fescue seed (*P* = 0.003, Table 1). On average, aNDF disappearance of endophyte-infected seed was lower than that of endophyte-free seed (11.53 ± 0.97% vs. 14.02 ± 0.96%, respectively). Additionally, isoflavone treatment significantly affected aNDF disappearance (*P* = 0.0053), where isoflavone powder treatment resulted in 14.28 ± 0.93% disappearance, and isoflavone tablet treatment resulted in 13.21 ± 0.93% disappearance, whereas control treatment resulted in 10.82 ± 0.93% disappearance. There was no difference (*P* = 0.49) in aNDF disappearance between powder or tablet forms of isoflavone.

**Table 1.** Effects of isoflavones and fescue seed type on neutral detergent fiber (aNDF), acid detergent fiber (ADF) and crude protein (CP) disappearance after rumen fluid incubation (48 h)

| Measurement<br>(%) | Isoflavone Treatment<sup>1</sup> |  |  |  |  |  | SEM | *P*-value<sup>3</sup> |  |  |
| --- | --- | --- | --- | --- | --- | --- | --- | --- | --- | --- |
|  | Control |  | Powder |  | Tablet |  |  |  |  |  |
|  | Fescue Seed<sup>2</sup> |  |  |  |  |  |  | Seed | Isoflavone<br>Treatment | Seed ×<br>Isoflavone<br>Treatment |
|  | E+ | E- | E+ | E- | E+ | E- |  |  |  |  |
| aNDF | 9.11 | 12.52 | 12.72 | 15.84 | 12.74 | 13.68 | 1.24 | 0.003 | 0.005 | 0.432 |
| ADF | 1.72<sup>c</sup> | 3.21<sup>bc</sup> | 7.01<sup>a</sup> | 2.19<sup>c</sup> | 1.27<sup>c</sup> | 5.91<sup>b</sup> | 1.03 | 0.36 | 0.009 | < 0.001 |
| CP | 2.44<sup>d</sup> | 12.23<sup>a</sup> | 6.98<sup>bc</sup> | 7.33<sup>bc</sup> | 4.01<sup>cd</sup> | 9.08<sup>ab</sup> | 0.93 | < 0.001 | 0.689 | 0.0002 |

<sup>1</sup>Treatment: control, isoflavone powder, or isoflavone pulverized tablet
<sup>2</sup>Seed type: KY 31 endophyte-infected (E+) tall fescue seed, KY 32 endophyte-free (E-) tall fescue seed
<sup>3</sup>Within a row, means among treatment groups are considered significant at $P \leq 0.05$ , and trending toward significance at $P \leq 0.10$
<sup>abcd</sup> Significant differences represented by means separation letters

ADF disappearance was associated with seed x treatment type interaction (*P* ≤ 0.05, Table 1). Effects of seed type on ADF disappearance was dependent upon isoflavone treatment (Table 1), as evident through a seed type by isoflavone treatment interaction (*P* < 0.0001). Samples that received the powdered isoflavone had greater (*P* = 0.0007) ADF disappearance from endophyte-infected tall fescue seed (7.01 ± 1.04 %) when compared to endophyte-free tall fescue seed (2.19 ± 1.04%). However, samples that received the tablet isoflavones had lesser (*P* = 0.001) ADF disappearance from endophyte-infected tall fescue seed (1.27 ± 1.04%) when compared to endophyte-free tall fescue seed (5.91 ± 1.04%). There were no statistical differences observed between powder and tablet (*P* = 0.23), nor between control and tablet (*P* = 0.18) on ADF disappearance.

There was a seed x treatment interaction on overall CP disappearance *(P* = 0.0002, Table 1). In the control isoflavone group, reduced CP disappearance was observed with endophyte-infected tall fescue seed type (2.44 ± 0.93 %) when compared to endophyte-free fescue seed (12.23 ± 0.93 %, *P* < 0.0001). There was no CP disappearance difference between endophyte-infected and endophyte-free seed when the powder isoflavone was used (*P* = 0.99); however, when the tablet isoflavone form was supplemented, endophyte-infected tall fescue seed yielded decreased CP disappearance (4.01 ± 0.99 %) compared to endophyte free tall fescue seed (9.08 ± 0.93 %, *P* = 0.01, Table 1).

### Ruminal VFA concentrations, bacterial populations, and pH

Propionate concentrations were significantly reduced for endophyte-free tall fescue seed (7.18 ± 0.63) when compared to endophyte-infected seed (5.71 ± 0.66, *P* = 0.03; Table 2), but no effect of treatment or treatment x seed interaction was observed (*P* > 0.1). The acetate: propionate ratio had a tendency to be reduced (*P* = 0.09; Table 2) with endophyte-infected tall fescue seed (1.74 ± 0.15) when compared to endophyte-free tall fescue seed (1.98 ± 0.15). No significant differences were observed among acetate, butyrate, valerate or IVMB concentrations (*P* > 0.1).

**Table 2.** Effects of isoflavones and fescue seed type on volatile fatty acid concentrations in the fluid phase of an *in vitro* rumen fluid digestion (48 h)

| VFA<sup>1</sup> | Isoflavone Treatment<sup>2</sup> |  |  |  |  |  | SEM | *P*-value<sup>4</sup> |  |  |
| --- | --- | --- | --- | --- | --- | --- | --- | --- | --- | --- |
|  | Control |  | Powder |  | Tablet |  |  |  |  |  |
|  | Fescue Seed<sup>3</sup> |  |  |  |  |  |  |  |  |  |
|  | E+ | E- | E+ | E- | E+ | E- |  | Seed | Isoflavone Treatment | Seed × Isoflavone Treatment |
| Acetate | 12.86 | 11.83 | 11.44 | 10.44 | 12.23 | 10.27 | 1.55 | 0.14 | 0.46 | 0.91 |
| Propionate | 7.96 | 6.09 | 6.00 | 5.52 | 7.57 | 5.52 | 0.95 | 0.03 | 0.31 | 0.64 |
| Butyrate | 2.00 | 1.65 | 2.00 | 1.44 | 1.73 | 1.64 | 0.43 | 0.18 | 0.90 | 0.81 |
| IVMB | 0.24 | 0.28 | 0.18 | 0.12 | 0.22 | 0.21 | 0.33 | 0.99 | 0.97 | 0.98 |
| Valerate | 0.36 | 0.15 | 0.21 | 0.21 | 0.27 | 0.26 | 0.11 | 0.21 | 0.69 | 0.29 |
| A:P | 1.60 | 1.96 | 1.91 | 1.99 | 1.73 | 1.98 | 0.21 | 0.09 | 0.62 | 0.73 |

<sup>1</sup> Concentration in mmol x L<sup>-1</sup>
<sup>2</sup> Treatment: control, isoflavone powder, or isoflavone pulverized tablet
<sup>3</sup>Seed type: KY 31 endophyte-infected (E+) tall fescue seed, KY 32 endophyte-free (E-) tall fescue seed
<sup>4</sup>Within a row, differences between treatment groups are considered significant at $P \leq 0.05$ , trending toward significance at $P \leq 0.10$
<sup>abcd</sup> Significant differences represented by means separation letters

After stringent sequence processing, a total of 408,813 high quality reads were obtained and averaged 17,034 ± 3,180 reads per sample, which is a consistent depth for sequencing analyses from ruminal samples [33]. Number of observed OTUs totaled 48,867 and averaged 2,037 ± 169 per sample. PERMANOVA global pairwise comparisons indicated significant differences among treatment groups (*P* = 0.03, *R^2^* = 0.32), however Shannon’s Diversity Index was not affected by 2 Concentration in mmol × L^−1^ treatment group (*P* > 0.1, Fig 1). Non-metric multidimensional scaling (NMDS) was utilized to analyze beta diversity (Figs 2, 3), where clusters of samples represent similarity of 16S rRNA bacterial genera by group based on rank. While treatment groups may have appeared to cluster similarly, there was no observable pattern between groups. Relative proportions of the genus level diversity between groups are represented in Fig 4. No differences in taxonomic profile were observed among groups (*P* > 0.1). However, finer changes in OTU abundances were identified. Numerous taxa were significantly different by seed, treatment or seed x treatment interaction *(P* < 0. 05; Table in S1 Table).

**Fig 1.**
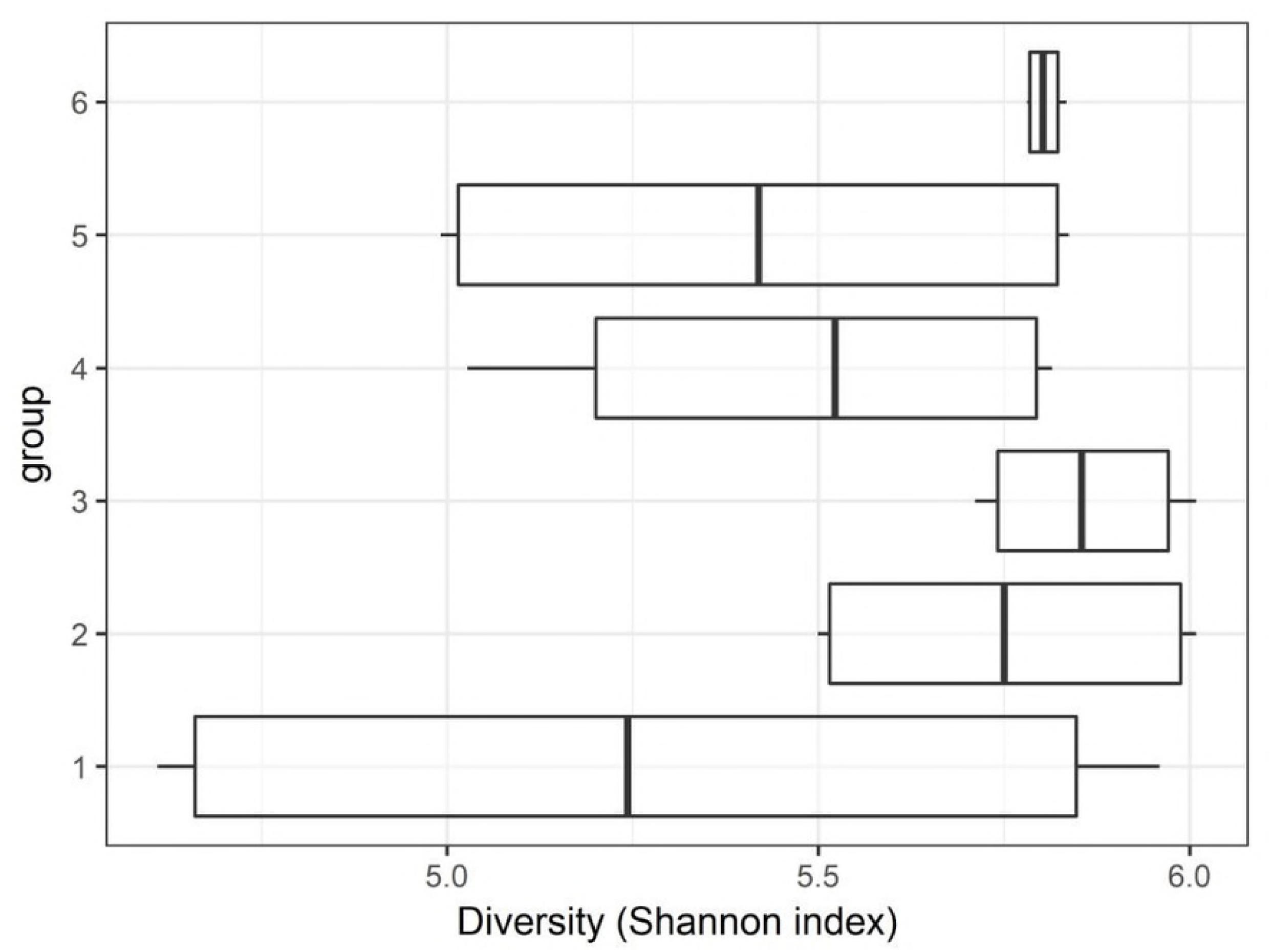
Shannon’s Diversity Index box plot of bacterial species diversity across treatment groups. Group 1: endophyte-infected tall fescue seed. Group 2: endophyte-free tall fescue seed. Group 3: endophyte-infected tall fescue seed with isoflavone powder. Group 4: endophyte-infected tall fescue seed with isoflavone tablet. Group 5: endophyte-free tall fescue seed with isoflavone powder. Group 6: endophyte-free tall fescue seed with isoflavone tablet.

**Fig 2.**
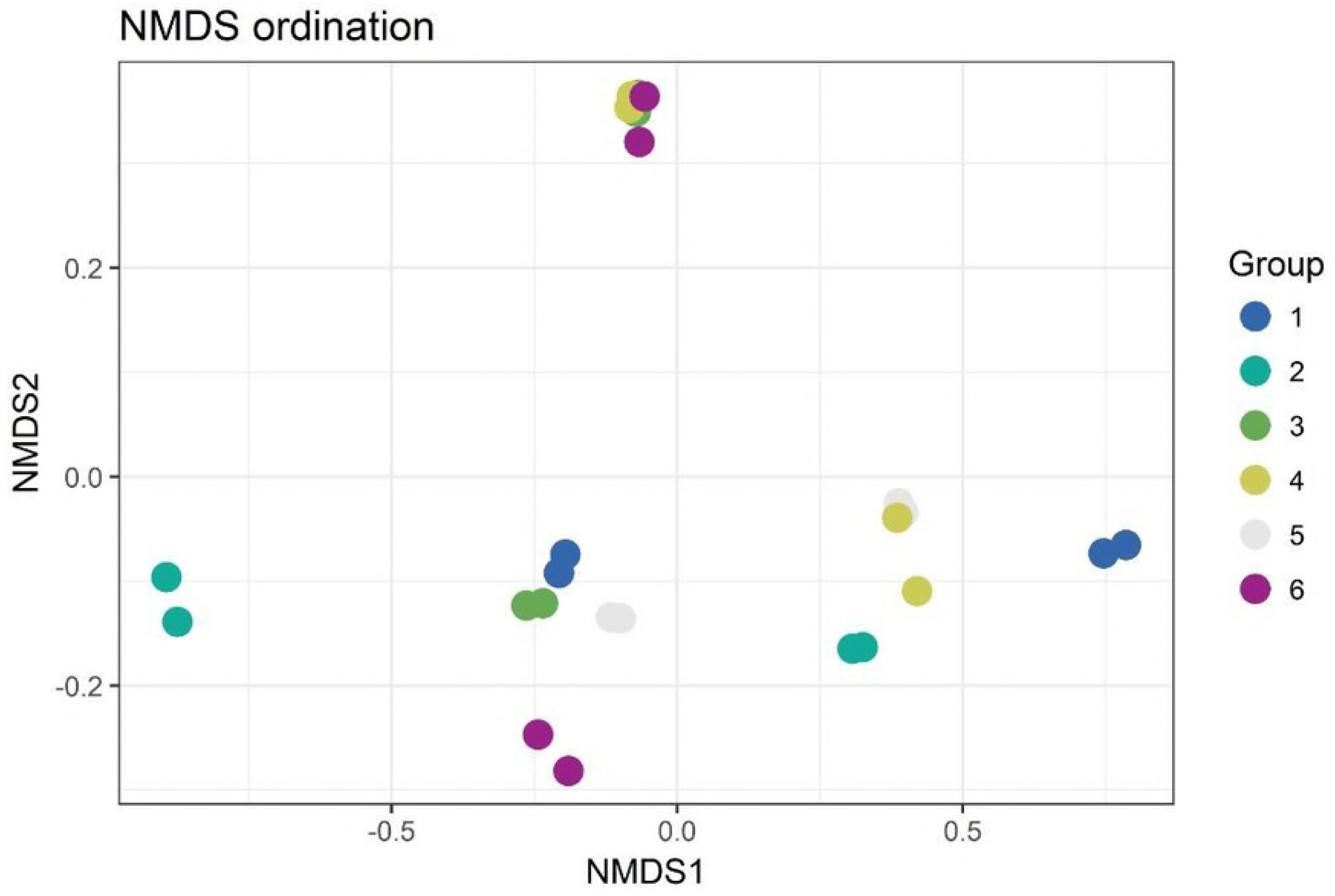
NMDS ordination plot grouped by treatment. Points represent taxa within treatment groups. Similar clustering indicates similar taxa within treatment groups. Group 1: endophyte-infected tall fescue seed. Group 2: endophyte-free tall fescue seed. Group 3: endophyte-infected tall fescue seed with isoflavone powder. Group 4: endophyte-infected tall fescue seed with isoflavone tablet. Group 5: endophyte-free tall fescue seed with isoflavone powder. Group 6: endophyte-free tall fescue seed with isoflavone tablet.

**Fig 3.**
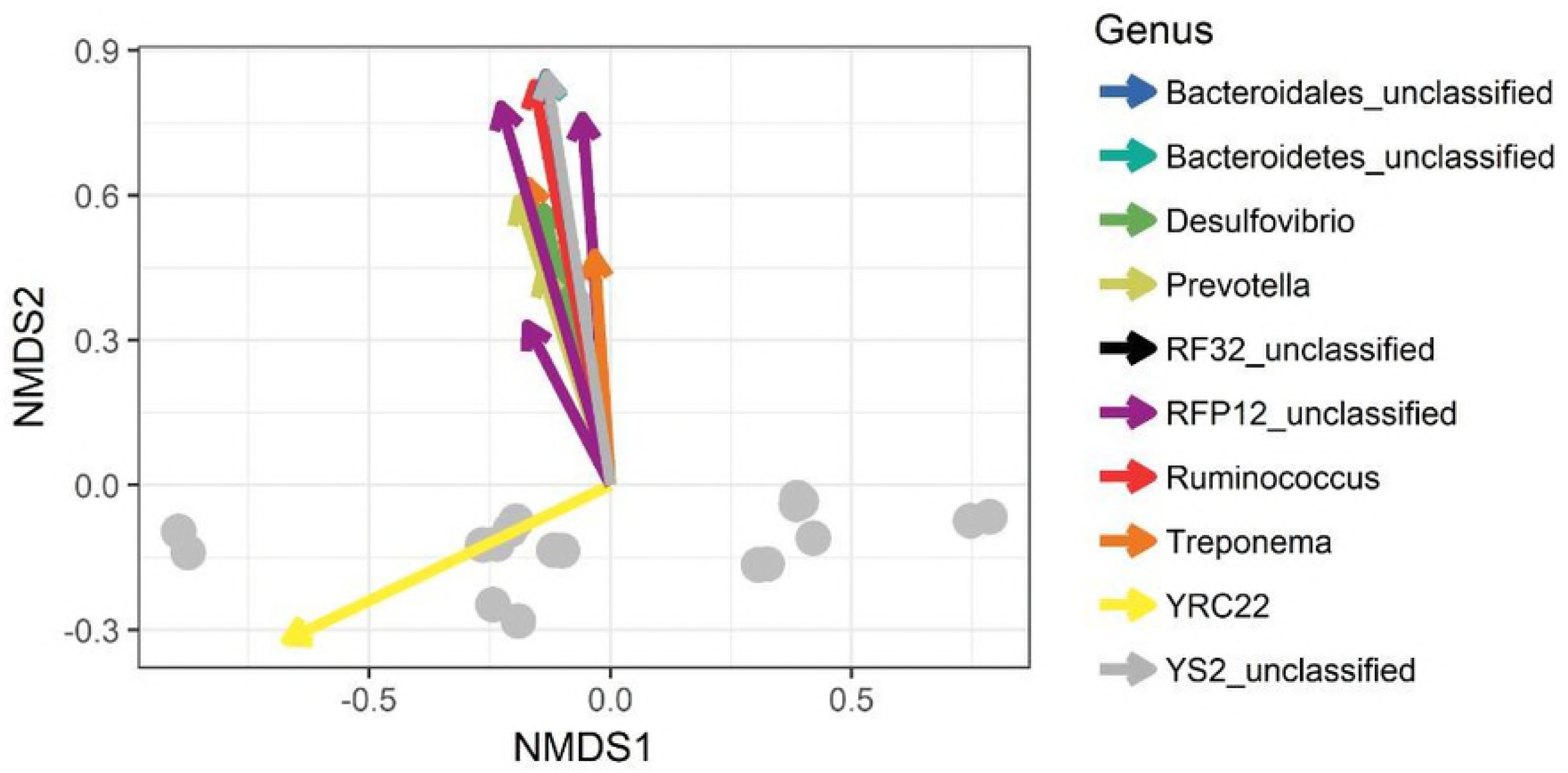
NMDS ordination plot indicating significant taxa at the genus level that influenced major bacterial shifts among treatments.

**Fig 4.**
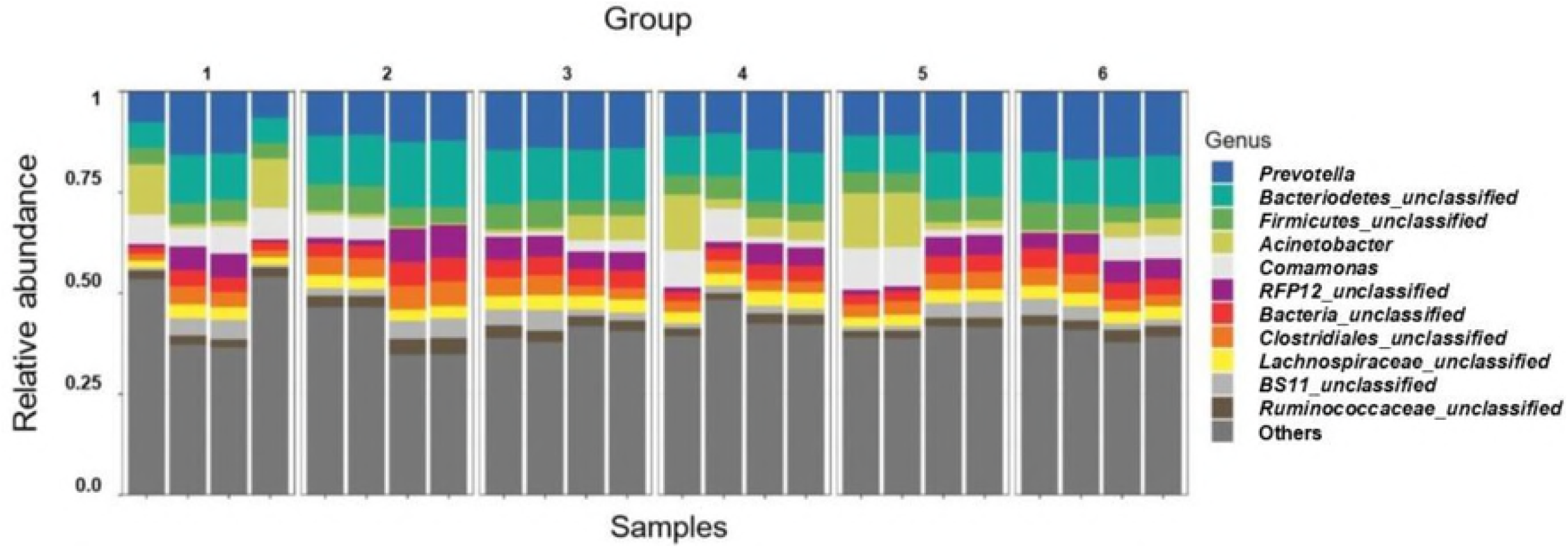
Taxonomic profiles of the relative proportions of bacterial communities by genus, grouped by treatment. Group 1: endophyte-infected tall fescue seed. Group 2: endophyte-free tall fescue seed. Group 3: endophyte-infected tall fescue seed with isoflavone powder. Group 4: endophyte-infected tall fescue seed with isoflavone tablet. Group 5: endophyte-free tall fescue seed with isoflavone powder. Group 6: endophyte-free tall fescue seed with isoflavone tablet. No significant differences among groups (*P* > 0.1).

Rumen pH was not significantly affected by seed type, treatment or seed x treatment interaction (*P* > 0.1; Fig 5), and pH ranged from 6.97 to 7.57.

**Fig 5.**
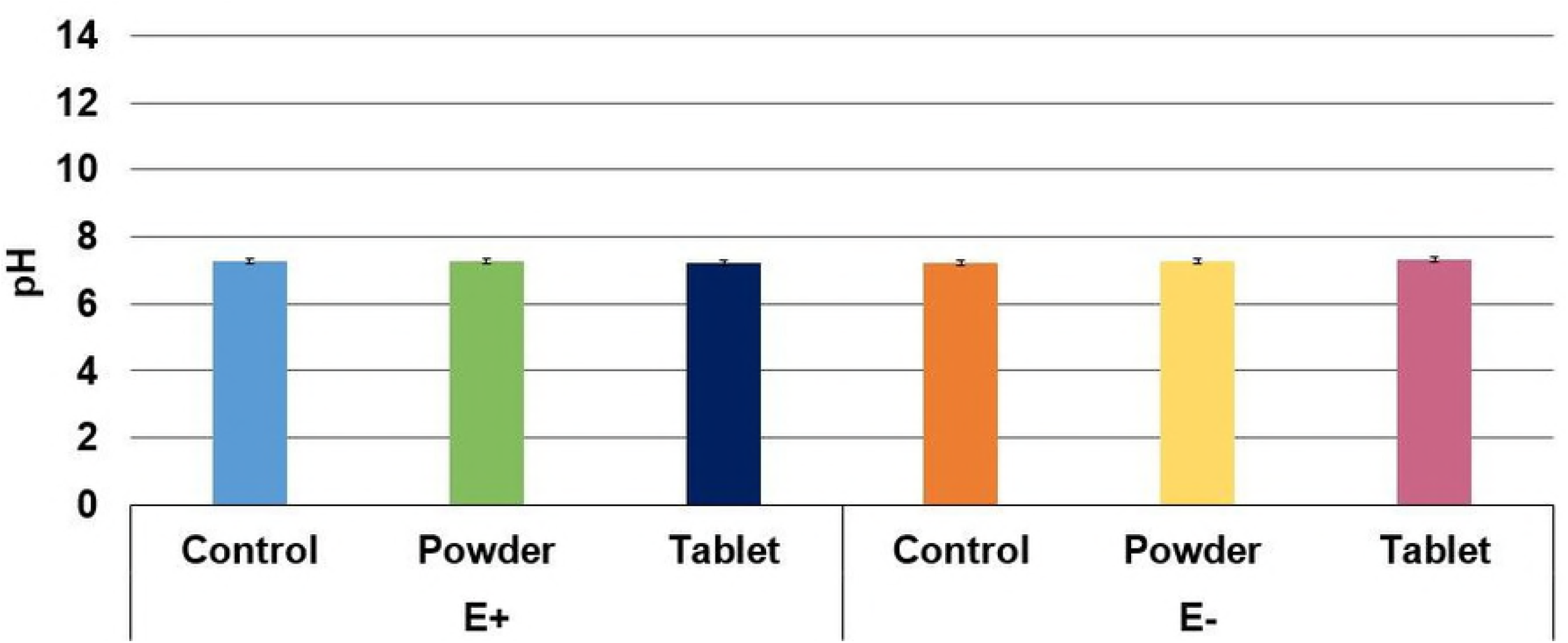
Mean ruminal pH by endophyte-infected (E+) and endophyte free (E-) seed type and treatment type. No significant differences in ruminal pH by seed, treatment or seed x treatment (*P* > 0.1). Error bars represent SEM.

## Discussion

Previous research conducted by Flythe and Kagan (34) and Harlow et al. [23] that noted improved fiber degradation and ammonia reduction employed biochanin A as the primary isoflavone (30 mg × L^−1^). The present study observed similar improvements in fiber degradation, including only 920 μg biochanin A per fiber bag (0.46 mg × L^−1^) and a total isoflavone concentration of 1.10 mg × L^−1^. The results from this experiment were consistent with those achieved by Harlow et al (2018), however the amount of total isoflavones (1.10 mg L^−1^) in this experiment may have been insufficient to elucidate all of the beneficial effects on fiber degradation and ammonia reduction that were previously reported.

The increase in ADF disappearance observed in the present study is consistent with what Harlow (23) observed, when the inclusion of the isoflavone biochanin A reduced overall ADF values post fermentation compared to the controls. As acid detergent fiber measures the amount of cellulose, which is a relatively indigestible fiber component, the inclusion of isoflavones with tall fescue seed in the present study increased indigestible fiber disappearance. ADF has an inverse relationship with digestible energy, and thus a high ADF component indicates less digestible energy available to the animal. In the present study, isoflavones resulted in a higher ADF disappearance, indicating possible improvement in available energy from tall fescue seed.

The results of CP disappearance in the present study are inconsistent with most tall fescue and CP literature, where proteolytic bacteria including several species of hyper-ammonia producing bacteria (HAB) are more readily able to degrade ergot alkaloid compounds [19] such as those found in endophyte-infected tall fescue. While these hyper-ammonia bacteria are able to degrade ergovaline, they are a source of inefficiency in the animal when dietary nitrogen is lost as ammonia. Antimicrobials, including ionophores and similar compounds, may be employed in the diet to combat this loss of nitrogen, selectively inhibiting some of the microflora and reducing wasteful byproducts of digestion [35–37]. Flythe et al. [34, 38] examined red clover isoflavones as antimicrobials due to their resistance to amino-acid degradation and action on HAB. They determined that biochanin A inhibited growth of *Clostridium stricklandii,* a member of HAB, and may prevent amino acid fermentation. A seed x treatment interaction affected CP disappearance and in general, increased disappearance was observed when endophyte-free tall fescue seed was fed while either no isoflavones were supplemented or when isoflavone was given in tablet form. However the isoflavone concentration used in the present study may not have provided a sufficient amount of isoflavones to elicit a reduction in ruminal amino acid degradation.

The donor animals (n = 2) used for rumen fluid procurement were not tall fescue-naïve animals, and consumed a forage-based diet. Rumen fluid obtained for this experiment may have been previously exposed to ergot alkaloids, which may have affected the results of ruminal pH and VFA concentrations. Additionally, ruminal microbial populations may have adapted or shifted from previous exposure to ergot alkaloids. The lack of significant differences in overall VFA concentrations (Table 2) provides support that the treatments did not inhibit or improve *in vitro* rumen fermentation. Richards, Pugh (39) similarly described no differences in rumen pH or VFA concentrations when steers consumed endophyte-infected tall fescue with or without soybean hulls. Rumen pH samples were not below 6.0 across replications of the experiment indicating fiber digestion and cellulolytic bacteria function [40] was not reduced.

Fiber degradability and VFA concentrations were notably reduced in the study. This was due to the fermentable substrate, as this study aimed to interrogate the effects of isoflavone supplementation on endophyte-infected tall fescue seed fermentation. Other studies have noted the benefits of utilizing ground tall fescue seed as a means of specifically examining the effects of ergot alkaloid exposure [41–44]. Consistent with the previous studies, the current experiment reflects similar biologically significant results, including those regarding fiber degradation (Flythe and Kagan, 2010; Harlow et al. 2018), which support the conclusion that isoflavones may be effective at improving fiber utilization when supplemented. Although VFA concentrations were unexpectedly lower, several studies using *in vitro* fermentation have noted reduced concentrations using similar methodology (Flythe and Aiken, 2010) compared to what is observed in the animal and subsequent studies should be performed.

Overall, 36 taxa were significantly different by seed, treatment or seed x treatment interaction (Table in S1 Table). Studies by Harlow (22) observed several species of amylolytic that may be inhibited with biochanin A: *Strep. bovis JB1, Strep. bovis HC5, Lactobacillus reuteri, Selenemonas ruminatium.* Additionally Harlow et al. [23] indicated several species of cellulolytic bacteria that are sensitive to biochanin A: *Fibrobacter succinogenes S85, Ruminococcus flavefaciens 8 and Ruminococcus albus 8,* however the same species shifts that occurred in studies conducted by Harlow et al. were not observed in the present study. Rather than being inhibited, *Selenemonas* proportions were increased when supplemented with isoflavones. However, in both studies conducted by Harlow et al., biochanin A was the only isoflavone supplemented and thus subsequent studies should be completed to verify population shifts with other isoflavones.

## Conclusions

The combined results of this study indicate some moderate rumen fermentation changes may have occurred due to treatments, but these results may have been affected by several factors. Where both fiber and crude protein disappearance results were not consistent with previous *in vitro* studies, this may be the result of a lower level of isoflavone administration to the *in vitro* fermentation system. As previous studies utilized a minimum of 30 mg × L^−1^ of biochanin A to elucidate responses, the present study utilized 0.63 mg × L^−1^ of biochanin A, which may have not been enough to elicit a response. Ergot alkaloid pressure on rumen fermentation was not clearly elucidated in the present study. VFA concentrations are indicative of rumen fermentation, which was not affected by each treatment. It is noteworthy that isoflavone administration did not alter VFA production, and that seed type was the driving factor in any of the changes in VFA concentrations. Foote et al. [45] noted significant reductions in VFA flux and absorption in animals administered endophyte-infected tall fescue seed, lending support that nutrient utilization and energy absorption may be further delayed when cattle experience tall fescue toxicosis. As propionate supplies up to 65% of the glucose carbon in cattle [46], and livestock consuming endophyte-infected tall fescue produce lower levels of VFAs [43], it is interesting that propionate was reduced by treatment with endophyte-infected seed in the present study. Foote, Kristensen (45) reported similarly reduced propionate concentrations, when rumen buffers containing extracts of ergovaline were applied to heat stressed ruminally cannulated steers. This reduction in propionate concentrations observed in the present study, as well as reduced total VFAs described by Foote, Kristensen (45), may provide further evidence that consumption of endophyte-infected tall fescue may reduce nutrient absorption.

While rumen bacterial populations varied among groups, the rumen microbiome is a dynamic ecosystem that is affected by various environmental and dietary factors, and thus changes in the bacterial taxa among groups may not be solely indicative of treatment with fescue seed or isoflavone treatment. As other studies have noted specific bacterial species capable of ergovaline degradation and benefits from isoflavones, further research should be conducted to validate consistent shifts in the rumen microbiome from ergot alkaloid pressure. Further research should be conducted to determine overall livestock, and more specifically cattle, performance with high ergot alkaloid pressure and mitigation of fescue toxicosis using isoflavones dosed at different concentrations.

## Acknowledgements

The authors would like to thank Gloria Gellin, USDA-ARS, for HPLC analyses.

## Supporting Information

S1 Table. Effects of isoflavones and fescue seed type on relative abundance of significant in vitro rumen bacterial taxa

